# Joint subject-identification and task-decoding from inferred functional brain graphs via a multi-task neural network

**DOI:** 10.1101/2023.11.27.568799

**Authors:** Elif Sema Balcioglu, Berkay Doner, Ekansh Sareen, Dimitri Van De Ville, Hamid Behjat

## Abstract

Functional connectivity (FC) between brain regions as manifested via fMRI entails signatures that can be used to differentiate individuals and decode cognitive tasks. In this work, we use methods from graph structure inference to estimate FC, which is in contrast to the conventional approach of deriving FC via correlation. Moreover, we infer FC graphs from seed-based co-activation patterns instead of raw fMRI data. We also propose a multi-task neural network architecture to jointly perform subject-identification and task-decoding from inferred functional brain graphs. We validate the developed model on data from the Human Connectome Project across eight fMRI tasks. Most importantly, our results show the superior task-decoding performance of FC graphs inferred from seed-based activity maps over graphs inferred from raw fMRI data. Furthermore, via gradient-based back-projection, we derive a significance score for inputs to the neural network, and present results showing the differential role of brain connections in subject-identification and task-decoding.

## 1. INTRODUCTION

An increasing number of studies have demonstrated the effectiveness of functional connectivity (FC) as a fingerprint for identifying individuals from a population (subject-identification) [1] and the cognitive tasks they perform (task-decoding) [2]. Conventionally, FC is estimated via computing the correlation between time-courses of pairs of brain regions. Alternatively, functional connectivity estimation via learning-based graph structure inference [3, 4] has shown promising results on both fMRI [5] and EEG [6, 7] data. The key objective of this work is to validate the added predictability power of FC graphs inferred from seed-connectivity maps over FC graphs inferred from raw fMRI data as in prior work. The secondary objective of this work is to present a novel calssification architecture that jointly performs subject-identification and task-decoding using a single trained model. Subject-identification is conventionally performed via building an identifiability matrix via correlations between test and re-test FC graphs [8, 9, 10], whereas for task-decoding, using support vector machines [11, 5] is the more common approach. In this study, we leverage the power of neural networks and develop a multi-task neural network (MTNN) model, which optimizes for multiple losses simultaneously, offering advantages such as enhanced generalizability and reduced overfitting [12]. We assess the robustness and reliability of graphs inferred from seed-connectivity maps using the MTNN within the context of multiple experiments.

## 2. METHODS

## 2.1. Functional learned graphs

Let 𝒢 denote an undirected, weighted graph with *N* vertices, and an edge set *ε* ⊂ {(*i, j*) | *i* ≠ *j, i, j* ∈ {1,…,*N*}}. Let **A** denote the *N* × *N* adjacency matrix of 𝒢, with elements *A*_*i,j*_ ∈ ℝ^+^ denoting the weight of edge (*i, j*). Let **D** denote the degree matrix of 𝒢, a diagonal matrix with elements *D*_*i,i*_ = Σ_*j*_ *A*_*i,j*_ . The combinatorial Laplacian matrix of *G* is given as **L** = **D** – **A**. Let **f** ∈ ℝ^*N*^ denote a graph signal whose i-th component **f** [*i*] is the signal value at the *i*-th vertex of *G*. The variability of **f** can be quantified using a measure of total variation (TV) as [13]:

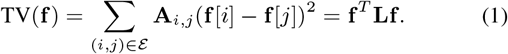

Given two signals **f**_1_ and **f**_2_, if TV(**f**_1_) < TV(**f**_2_), then **f**_1_ is smoother than **f**_2_ as it exhibits less variability on *G*. Using this notion of smoothness, a sparse graph structure can be inferred from a given set of observations [3]. Let **F** denote an *N* × *M* matrix where each column is an observation, a graph signal, and **Z** an *N* × *N* matrix with elements *Z*_*i,j*_ = ∥**F**_*i*,:_ – **F**_*j*,:_∥_2_, i.e., Euclidean distance between signal values on vertices *i* and *j*. A graph structure can be inferred from **F** via the optimization [4]:

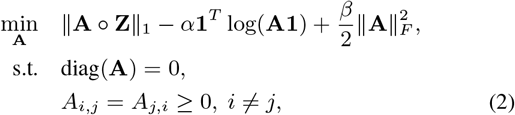

where *o* is the Hadamard product, α and β are regularization parameters, ∥*·*∥_*F*_ the Frobenius norm, and **1** = [1,. .., 1]^*T*^ . The first term in (2) enforces smoothness by invoking that [4]:

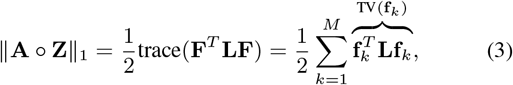

Smooth graph signals reside on strongly-connected vertices, thus, the vertices are expected to have smaller distances. The second term enforces degrees to be positive and improves overall connectivity.

### 2.2 Multi-task neural network

With a sample size limited to 100 individuals and a relatively high number of input features (specifically, upper triangular entries of the adjacency matrix, i.e., *M* = *P* (*P* – 1)*/*2 features), our objective with using the MTNN is to efficiently share the weight-heavy components of the model, which maps the *M* input features to a lower dimensional representation of size *L* | *L* << *M* . Despite reducing the parameter size, as the initial representations are shared by two objectives, comparable performance can be achieved with other classification methods. The MTNN architecture produces two outputs, mapping the input features to both subject-identification and task-decoding labels. The input to the model is the upper triangular parts of the adjacency matrix of an FC graph. These flattened features are then mapped to an intermediate representation, which splits into two branches, one for subject-identification and one for task-decoding. In each step, we use fully-connected layers, followed by ReLU activation [14], and batch normalization [15] layers. We train the network using the cross-entropy losses of subject-identification (SI) and task-decoding (TD) problems. Specifically, given *S* subjects and *T* fMRI tasks, we train the model using a cross-entropy loss given as:

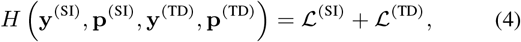

where ℒ^(SI)^ and ℒ^(TD)^ denote the cross entropy terms for SI and TD, respectively, defined as:

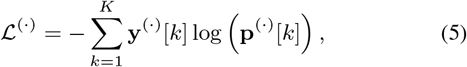

where **y**^(*·*)^[*k*] ∈ {0, 1} is ground truth label for class *k*, **p**^(*·*)^[*k*] ∈ (0, 1) is the softmax probability of the output for class *k*, and *K* is the number of classes. In our experiments, for subject-identification we had *K* = 100 subjects and for task-decoding we had *K* = 8 tasks. Fig. 1 shows our proposed MTNN architecture.

**Fig. 1.**
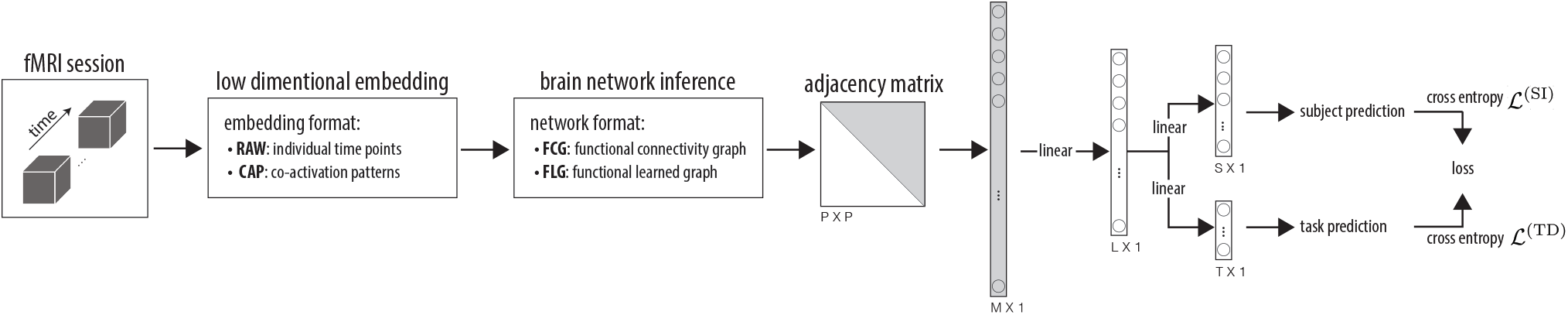
Pipeline for joint subject-identification and task-decoding from fMRI.

### 2.3 Experimental dataset

We obtained structural MRI and fMRI data from the group of 100 unrelated subjects (54% female, mean age = 29.11*±* 3.67, age range = 22-36) of the Human Connectome Project (HCP) dataset [16]. We used the HCP minimally preprocessed data [17], which ensures the necessary co-registration accuracy to apply this method. The fMRI data consisted of two repetitions of seven task-based sessions and a resting-state session per subject, thus, in total 16 sessions per subject. We used the minimally preprocessed volumetric fMRI data as provided by HCP. As for temporal preprocessing, for each voxel, we regressed out head movement, performed de-trending (0, 1st, and 2nd order), and, finally, z-scored across time. For each subject and each fMRI session, we transformed the high-resolution data (3D volumes) into two different low-dimensional embeddings. For both embeddings, we use the Schaefer brain atlas [18] with *P* = 400 parcels, each being a region of the cerebral cortex—with 200 parcels in each of the hemispheres; see Fig. 2(a).

**Fig. 2.**
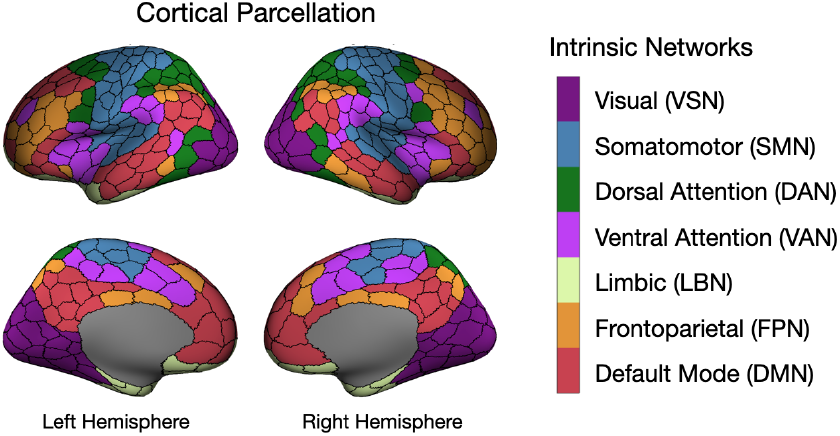
The Schaefer400 brain atlas [18]; each region is assigned to one of the seven intrinsic brain networks [19].

For the first embedding—denoted in the following as the `RAW` data—each fMRI frame (each frame representing one time point) was parcellated using the atlas, and, subsequently voxels within each parcel were averaged, resulting in *P* ×*K* matrices, where *K* denotes the length of the session. For the second embedding—denoted in the following as the `CAP` data—we extracted *P* static seed-based coactivation patterns, each associated with one atlas region. Each connectivity map was defined based on the method initially introduced in [20], and more recently used in [21]. Specifically, we extracted the average activity time course of each brain region, and then selected the top 15% of the frames (time-points) that showed maximal activity. These frames were then averaged, resulting in one 3D volume (a co-activation map) per brain region. Each co-activation map was then parcellated with the atlas, and voxels within each region were then averaged. By repeating this procedure across all atlas regions, each fMRI session was transformed to a *P* ×*P* matrix, each column representing a seed-connectivity map.

### 2.4 Experimental setups

Through a series of experiments, we used the MTNN architecture to compare the performances of FC matrices generated using data formats `RAW` and `CAP`, and methods functional learned graph (`FLG`) and functional connectivity graph (`FCG`). As a first experiment, we validated the performance of MTNN compared to two algorithms: Support Vector Machines (SVM) and Random Forest (RF). For this, we found the best parameters (*C*, γ for SVM with “rbf” kernel, maximum depth, and number of estimators for RF) with grid-search, using a 3-way cross-validation. It is crucial to note that MTNN consists of a single model predicting both subject-identification and task-decoding however, separate models for those objectives are trained for SVM and RF. For all experiments on MTNN, we used the Adam optimizer [22] with a 5e-4 learning rate and a 5e-4 weight decay.

Given the two repetitions available for each subject and each fMRI session type, we used the first repetition data for training and performed testing on the second repetition data. Specifically, we trained our models on 800 first repetition fMRI sessions (100 subjects × 8 task types), and subsequently tested the trained models on 800 second repetition sessions. This separation of the data was used in all our experiments, that is, for each data format (`RAW` or `CAP`) and each graph inference method (`FCG` or `FLG`).

In all our experiments on `FCG`, we used density-matched `FCG`, wherein we set the smallest edge weights to zero up to the point where the density of the matrix (i.e., number of remaining graph edges) becomes equal to that of `FLG` of the corresponding session. This enabled us to have a fair comparison between the two methods, as our initial experiments (results not presented here) revealed that fully-connected `FCG` perform substantially worse than `FLG` compared to density-matched `FCG`.

## 3. RESULTS AND DISCUSSION

In the first experiment we compared the performance of the MTNN on the `RAW` data against two widely used classification methods: SVM and Random Forest, trained separately for both objectives; see Table 1. The MTNN outperformed both SVM and Random Forest in task-decoding (TD) and exhibits a performance comparable to SVM in subject-identification (SI). Importantly, MTNN’s superior performance is obtained using a single trained model in contrast to the use of separate trained models for SI and TD in SVM and RF.

**Table 1.**
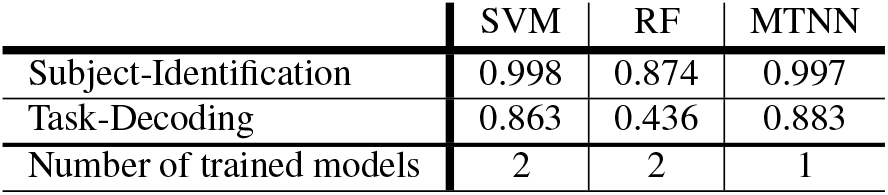
Accuracy of MTNN vs alternative methods.

|  | SVM | RF | MTNN |
| --- | --- | --- | --- |
| Subject-Identification | 0.998 | 0.874 | 0.997 |
| Task-Decoding | 0.863 | 0.436 | 0.883 |
| Number of trained models | 2 | 2 | 1 |

Fig. 3(a) shows subject-identification and task-decoding performances on `FCG` and `FLG` derived from the `RAW` data. The results are presented for varying numbers of frames, ranging from as few as 10 frames to 100 frames. With the exception of 10 frames, `FLG` outperforms `FCG` on SI, with accuracy reaching very close to 1.0 after 60 frames, whereas `FCG` remains at an accuracy of 95%. On TD, `FCG` catches up to the performance of `FLG` as we increase the number of frames, however, with only a few frames, `FLG` shows a substantial advantage, with around 60% accuracy even with as few frames as 10. Fig. 3(b) shows individual performances on eight different tasks on TD. The figure reports the accuracy difference between `FLG` and `FCG`.`FLG` shows superior performance over `FCG` across the majority of tasks and across different number of frames. In particular, `FLG` shows as much as 50% improved performance over `FCG` when a minimal number of frames are used.

**Fig. 3.**
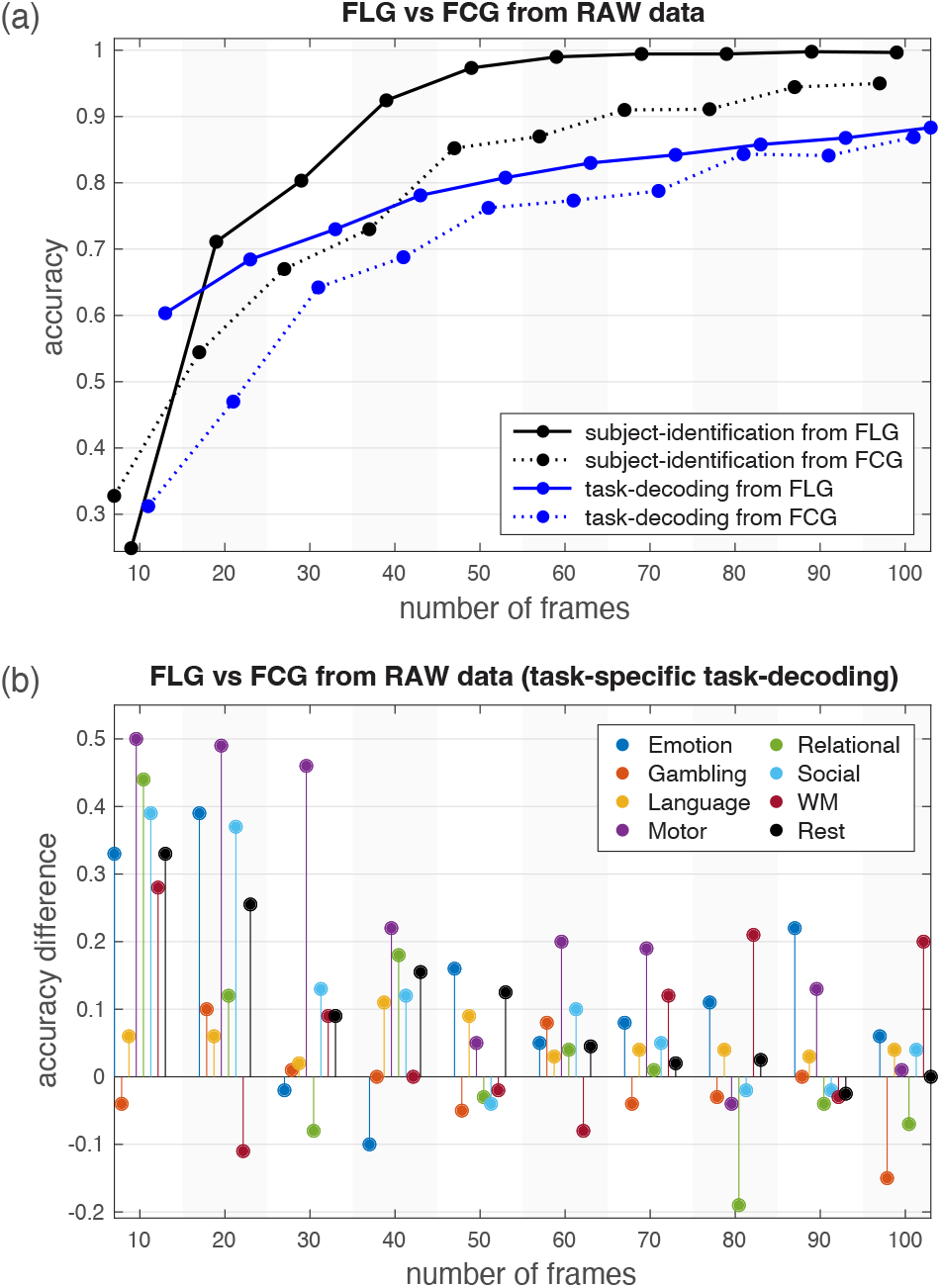
(a) SI and TD prediction accuracy on data `RAW` using `FCG` and `FLG` graphs. (b) Average across tasks TD accuracies shown in (a) are split into eight different tasks, and the difference in performance between `FLG` and `FCG` is reported; values greater than zero indicate superior performance of `FLG` over `FCG`, and vice versa.

We then assessed the performance of the `CAP` data. In addition to analyzing the data as a whole—i.e., using the entire *P* × *P* matrices as input for graph inference, we inferred graphs from seed-connectivity maps of regions that fall within each of the seven intrinsic brain networks—i.e., using only a subset of the columns from the data for each network, where the number of columns (regions) ranged from 26 (LBN) to 91 (DMN) depending on the network. Fig. 4(a) shows SI and TD performance differences of the FC derived from the `CAP` data using `FLG` and `FCG`. For each network, `FLG` consistently outperforms `FCG` in both SI and TD. We then aimed at assessing the SI and TD performance using `FLG` from the `RAW` and the `CAP` data. Since both formats have unique characteristics, it is difficult to find a setting where they are treated equally. As a solution, for a network with *R* regions, we generated a new data format based on the `RAW` data with the same shape as `CAP` (*P* × *R*), where each column was obtained by randomly selecting 15%—as used for generating the `CAP` data—of the columns from the `RAW` data and averaging them. We denote this version of the `RAW` data generated via random averaging of frames as `RAW-RAND`. Fig. 4(b) shows the accuracy difference `FLG` from `CAP` to `FLG` from `RAW-RAND`. `FLG` from the `CAP` data provide representations that enable superior TD across all brain networks. In contrast, for SI, `FLG` from the `RAW-RAND` data shows superior performance across all networks.

**Fig. 4.**
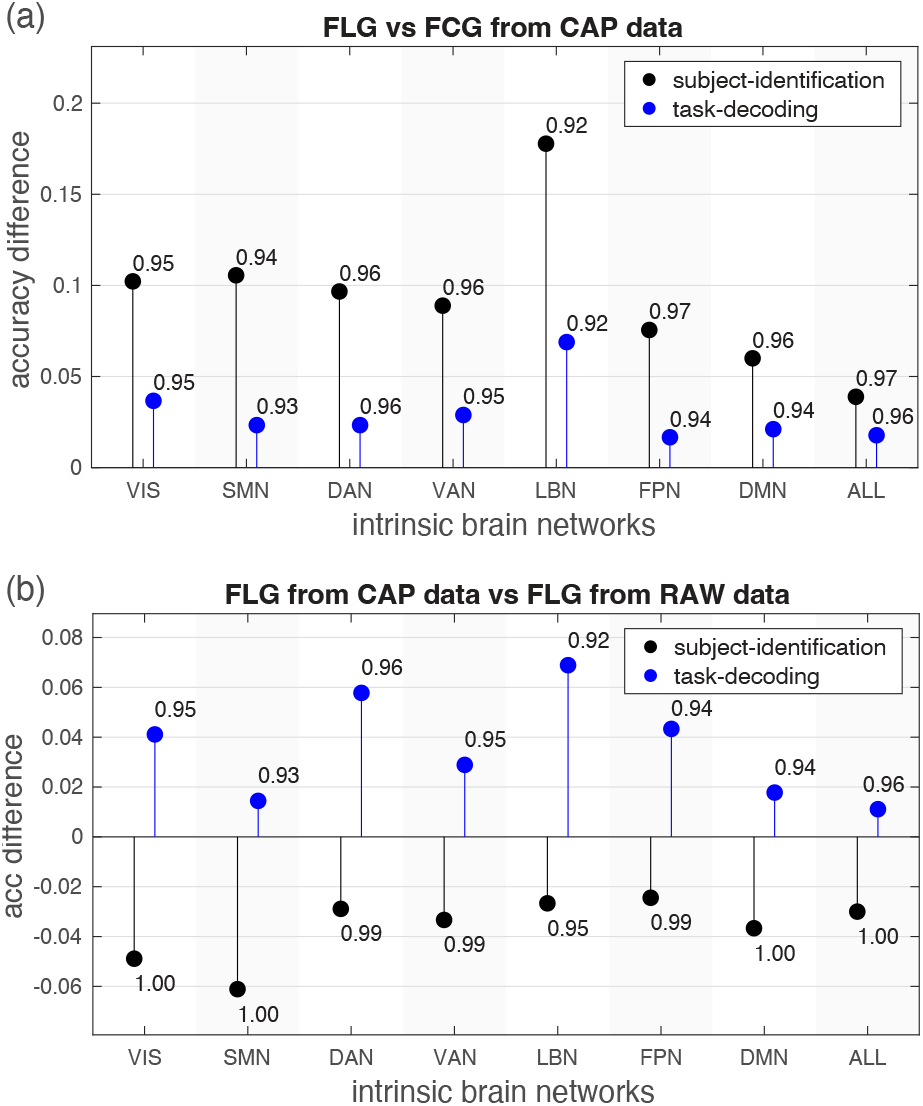
(a) SI and TD prediction accuracy on the `CAP` data using `FCG` and `FLG` for the seven intrinsic brain networks shown in Fig. 2. Values above zero indicate better performance using `FLG`, and vice versa. (b) SI and TD prediction accuracy using `FLG` graphs inferred from `CAP` data compared to `RAW` data. Values above zero indicate better performance on CAP data whereas value below zero indicate better performance on RAW data. In both (a) and (b), values at the tip of stem bars show the accuracy of the superior method.

To disentangle the discrepancy between the result on TD and SI of Fig. 4(b), we aimed at assessing the “significance” of input features to the networks—i.e., brain graph edge weights—on the resulting predictions. To this aim, we used the DeepLIFT algorithm [23], which is a gradient-based interpretability method. The significance of each edge is calculated from the difference in neuron activation values from a reference input. A reference input in our case refers to a functional connectome matrix consisting of zeros. The algorithm gives separate consideration to positive and negative contributions to the prediction, where values close to 0 show a lack of significance. Using this method, for each trained MTNN, we constructed two upper triangular *P* × *P* matrices, one for SI and one for TD, denoted **S**^(*TD*)^ and **S**^(*SI*)^, respectively. Each upper triangular element (*i, j*) of these matrices, denoted 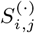, represents the significance of the associated graph edge—i.e., the connection between region *i* and region *j*—-in the success of the trained model for the task, i.e., TD or SI. Each element 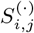 was obtained by averaging *T* or *S* values for TD and SI, respectively, each of which is the absolute value of per-class significance of edge (*i, j*). The averaging enables identifying edges that have the greatest significance across all classes. The values were then taken to the logarithm scale.

We constructed **S**^(*TD*)^ and **S**^(*SI*)^ for the MTNN trained on `FLG` from the `CAP`, denoted 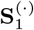, and also for that trained on `FLG` from the `RAW-RAND` data, denoted 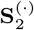. Fig. 5 shows the results. MTNN trained on `FLG` from the `CAP` data prioritize inter-network connections more than `RAW` for TD, shown in the left part of Fig. 5; note the large values found in the off-diagonal blocks relative to the block diagonal values. On the contrary, both networks give similar importance to intra-network connections, which can be seen in values close to zero on block diagonal segments. This observation on TD is similar for SI, as can be inferred from the right-hand plot, which shows the difference between the two prediction tasks (TD minus SI); TD on `FLG` from the `CAP` data in general shows lower edge significance than SI, as reflected by the negative values, despite differences being very small (see colorbar range). This similarity is not unexpected, given the shared representation of both objectives in the MTNN network. In contrast to `CAP`, `RAW-RAND` places greater weight on internetwork connections between the LBN and other networks (see dark blue regions), which may be a factor to explain the superior SI performance of `FLG` from the `RAW-RAND`.

**Fig. 5.**
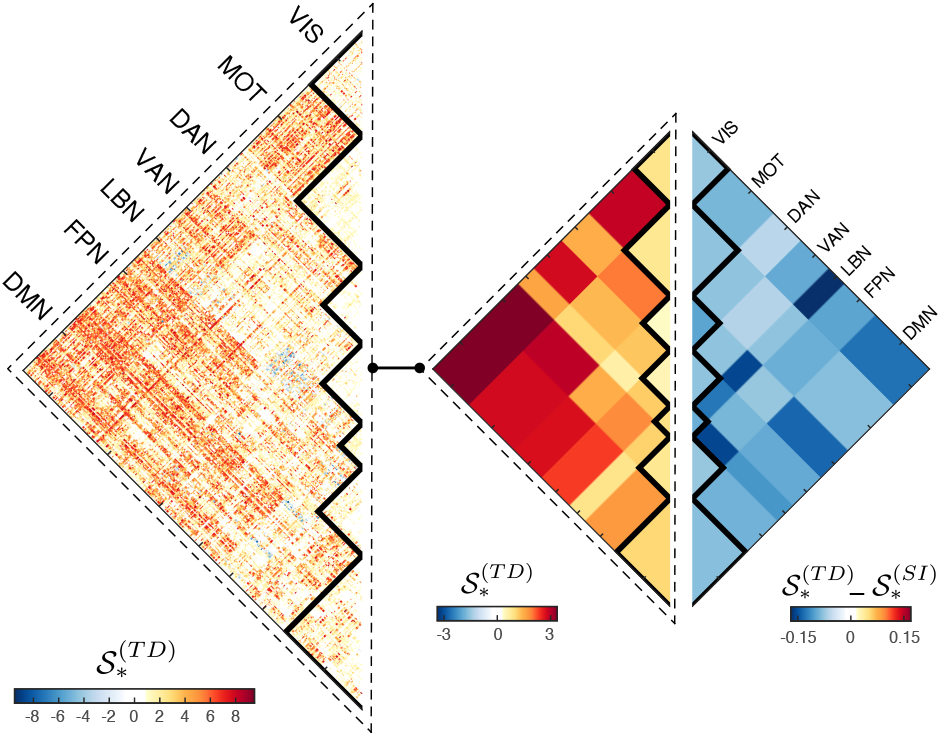
Difference in the significance of brain connections in task-decoding and subject-identification for MTNN trained on `FLG` from the `CAP` data and MTNN trained on `FLG` from the `RAW-RAND` data. Specifically, here we are assessing the trained models associated to the results reported in the last column (“ALL”) on Fig. 4(b).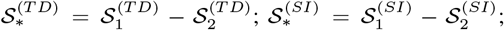 formal description of 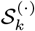 for *k* = 1, 2 is given in the main text. For the two half-matrices marked with dashed lines and shown as connected, the one on the right is obtained via computing the mean of the intra- and inter-network blocks of the one on the left, to ease visual comparison. The half matrix on the far right hand side was computed via the same method of block averaging.

## 4. CONCLUSIONS AND OUTLOOK

In this work we proposed a method for joint task-decoding and subject-identification from fMRI data. The proposed method initially embeds 4D fMRI data into a low dimensional space using a cortical atlas, specifically, constructing co-activation patterns linked to individual cortical regions [21]. A functional brain graph is then inferred from the low dimensional embedding [6], and, subsequently fed into the MTNN. The MTNN is trained to jointly predict the identity of the subject and the task being performed. Through a series of experiments, we show the superiority of the proposed method over conventional strategies of performing classification based on correlation-based FC. Our promising results suggest the potential of further expanding the proposed MTNN architecture, on different fronts: i) to take as input higher resolution FC graphs [24] or fMRI spectral features [11] extracted on macro-scale [25] or voxel-wise [26, 27] structural brain graphs; ii) to use graph neural networks [28] or methods of transfer learning to learn individualised task-decoding models [29]; iii) integrating the brain network inference step into the neural network. The present study only briefly touched upon disentangling the significance of different intra/inter network connections in task performance, an avenue which we believe is fertile to be explored in future work.

